# miRNAs, target genes expression and morphological analysis on the heart in gestational protein-restricted offspring

**DOI:** 10.1101/507038

**Authors:** Heloisa Balan Assalin, José Antonio Rocha Gontijo, Patrícia Aline Boer

## Abstract

**BACKGROUND:** Gestational protein restriction was associated with low birth weight, hypertension and higher prevalence of cardiac disorders in adults. Several mechanisms, including epigenetics, could be related with the cardiovascular phenotype on protein-restricted offspring. Thus, we investigated the morphological cardiac effects of gestational protein restriction and left ventricle miRNAs and target genes expression pattern in both 12-day and 16-week old gestational protein-restricted male offspring.

**METHODS:** Pregnant Wistar rats were allocated into two groups, according to protein supply during pregnancy: NP (normal protein diet-17%) or LP (low protein diet − 6%). The study evaluates the effects of maternal protein restriction on food consumption and body weight of both pregnant dams and offspring, systolic blood pressure in 16-wk old offspring and on cardiac morphometric and molecular parameters in both 12-d and 16-wk old offspring.

**RESULTS:** Dams on the gestational protein-restricted diet had lower body weight gain and higher food intake. Gestational protein-restricted offspring had low birth weight, followed by rapidly body weight recovery, hypertension, and increased myocytes crosssectional area and collagen fraction at 16-week old age. At 12-days old, miR-184, miR-192, miR-376c, miR-380-3p, miR-380-5p, miR-451, and miR-582-3p had increased expression, and miR-547 and miR-743 a had decreased expression in the gestational protein-restricted left ventricle. At 16-week old, let-7b, miR-125a-3p, miR-142-3p, miR-182 and miR-188-5p had increased expression and let-7g, miR-107, miR-127, miR-181a, miR-181c, miR-184, miR-324-5p, miR-383, miR-423-5p and miR-484 had decreased expression in gestational protein-restricted left ventricle. Target predicted gene expression analysis shown higher expression of Dnmt3a, Oxct1, Rictor and Trps1 and lower expression of Bbs1 and Calml3 in 12-day old protein-restricted offspring. 16-week old protein-restricted offspring had higher expression of Adrbk1, Bbs1, Dnmt3a, Gpr22, Inppl1, and Oxct1 genes.

**CONCLUSION:** Gestational protein restriction leads to offspring low birth weight, increased systolic blood pressure and morphological heart alterations that could be related to early heart miRNA expression changes that perpetuate into adulthood and which are associated with the regulation of essential genes involved in cardiovascular development, heart morphology, function, and metabolism.

## 1. INTRODUCTION

Several epidemiological and experimental studies have shown associations between gestational protein restriction, low birth weight and a higher prevalence of cardiovascular disease in adulthood [5,57]. Initially, it was thought that the mechanisms causing cardiovascular changes in protein-restricted offspring might be secondary to the development of arterial hypertension [41] and endocrine changes, such as insulin and leptin resistance [8,46]. Alternatively, taking into account the evidence, studies have shown that primary insults in heart development itself might predispose to cardiovascular dysfunction later in life. Thus, protein restriction in the intrauterine environment results in permanent changes in cardiac structure and function [4,49]. Several authors have shown maternal protein restriction to lead to impairment in offspring cardiomyocyte proliferation and differentiation [1,9], reduction of cardiomyocyte number [17,75], fibrosis [45,49] and, ultrastructural changes that may lead to impaired cardiac function later in life [17]. However, information regarding the molecular mechanisms of the etiopathogenesis of these cardiac changes is still scarce.

MicroRNAs (miRNAs) are genomic-encoded small noncoding RNAs of approximately 22 nucleotides in length that play an essential role in post-transcriptional regulation of target gene expression [2,6]. miRNAs control gene expression post-transcriptionally by regulating mRNA translation or stability in the cytoplasm [52]. Although only recently discovered [42], it has become clear that miRNAs are critical components of diverse regulatory networks in animals [2].

Functional studies indicate that miRNAs are involved in critical biological processes during development and in cell physiology [6,10], and changes in their expression are observed in several pathologies [10,13]. Currently, it is known that miRNAs are not only involved in cardiovascular development and physiology [12,60] but also in several cardiovascular diseases [56,67]. Therefore, the expressional study of miRNAs and target genes that undergo miRNA-mediated regulation in the heart may help the understanding of the mechanisms underlying the cardiovascular phenotype on protein-restricted offspring.

This study aimed to evaluate the miRNAs and predicted gene expression pattern of rat cardiac left ventricle (LV) in both 12-day and 16-week old gestational protein-restricted male offspring in an attempt to elucidate the possible molecular mechanisms involved with the etiology of the cardiac phenotype observed in gestational protein-restricted offspring. Furthermore, we wished to evaluate the effects of maternal protein restriction on food consumption and body weight of both pregnant dams and offspring, systolic blood pressure in 16-wk old offspring and on cardiac morphometric parameters in both 12-d and 16-wk old offspring.

## 2. METHODS

### 2.1 Experimental protocol

#### Animals and diets

The Institutional Ethics Committee (CEUA/UNICAMP #315) approved the experimental protocol, and the general research guidelines on animal care established by the Brazilian College of Animal Experimentation (COBEA) and by NIH Guide for the Care and Use of Laboratory Animals were followed throughout the investigation. The experiments were conducted as described in detail previously [49,50] on age-matched rats of 12-week-old sibling-mated *Wistar HanUnib* rats (250–300 g). The local colonies originated from a breeding stock supplied by the Multidisciplinary Center for Biological Investigation on Laboratory Animal Science, UNICAMP, Brazil. Male and female weanling *Wistar HanUnib* rats were housed and maintained under a 12-hour day/night cycle (lights on 06.00 – 19.00 h) at constant temperature (22±2°C), with standard chow (Nuvital, Curitiba, PR, Brazil) and water available *ad libitum*. From 12 to 14 weeks of age, the animals were mated. Pregnant dams were singly-caged and randomly assigned either the regular protein diet (NP, 17% casein) or isocaloric low protein diet (LP, 6%) throughout the entire pregnancy. Body weight and food intake were evaluated weekly in pregnant dams. Birth weight and anogenital distance were measured in male offspring. Litter size was adjusted at birth to eight. At 12 days after birth, half of the male offspring of each dam was euthanized (NP-12d and LP-12d groups). At 21 days after birth, the remaining male offspring were weaned and caged separately. Body weight was evaluated weekly from birth to 16 weeks after birth, and food intake was assessed daily from weaning to 16 weeks after birth when they were euthanized (NP-16w and LP-16w groups).

### 2.2 Blood Pressure Measurement

The systolic blood pressure was measured in conscious male offspring from 6 to 16 weeks after birth, employing an indirect tail plethysmography method. Briefly, an indirect tail-cuff method using an electro-sphygmomanometer combined with a pneumatic pulse transducer/amplifier was used (IITC Life Science—BpMonWin Monitor Version 1.33). Measurements were conducted at the same time during the day. This indirect approach allowed repeated measurements with close correlation (correlation coefficient = 0.975) compared with direct intra-arterial recording. The mean of three consecutive readings was taken as the blood pressure.

### 2.3 LV weight measurement

At 12 days and 16 weeks of age, some male offspring from different litters were deeply anesthetized with a mixture of ketamine (50 mg/kg body weight, *i.p*.) and xylazine (1 mg/kg body weight, *i.p*.). Heart LV was dissected, weighed and stored at −80°C. The tibia length was also measured.

### 2.4 Histological Analysis

At 12 days and 16 weeks of age, some male offspring from different litters were anesthetized and had the heart LV perfused with a heparinized saline solution (1%) and with a 4% (w/v) paraformaldehyde solution in 0.1M phosphate buffer (pH 7.4). After perfusion, the LV was dissected, fixed for 24 hours in the paraformaldehyde solution, and then embedded in paraplast (Sigma-Aldrich^®^). Five-micrometer-thick sections were stained with hematoxylin and eosin (HE) or picrosirius red. The measurements were performed from digital images that were collected by a video camera attached on an Olympus microscope (x40 magnification lens), and the images were analyzed by Image J software. The cross-sectional area (CSA) was measured with a digital pad; the selected cells were transversely cut so that the nucleus was in the center of the myocyte and, determined as an average of at least 30 myocytes per animal. The heart interstitial collagen volume fraction, marked by picrosirius, was calculated as the ratio between the connective tissue area and connective tissue plus myocyte areas, from 30 microscope fields of digitalized images of each animal. Perivascular collagen was excluded from analysis.

### 2.5 Heart LV miRNA Expression

Four male offspring from different litters were used in each group for the miRNA expression analysis. Total ribonucleic acid (RNA) was extracted from LV samples using Trizol reagent (Life Technologies, USA) [15]. Total RNA was quantified (Take3 micro-volume plate of the Epoch spectrophotometer; BioTek^®^, USA). The RNA integrity was evaluated by electrophoresis on a denaturing agarose gel stained with GelRed Nucleic Acid Gel Stain (Uniscience, USA) and the RNA purity was assessed by the ratio of absorbance at 260 and 280 nm. Briefly, 450 ng RNA was reverse transcribed using TaqMan^®^ MicroRNA Reverse Transcription Kit and Megaplex RT Primers Rodent Pool A (Life Technologies, USA), according to the manufacturer’s guidelines. Complementary DNA (cDNA) was amplified using a TaqMan^®^ Rodent MicroRNA Array A v2.0 with TaqMan Universal PCR Master Mix on QuantStudio 12K Flex System (Life Technologies, USA), according to the manufacturer’s instructions. Data analysis was performed using relative gene expression evaluated using the comparative quantification method^47^. The U87 gene was used as a reference gene. Mean relative quantity was calculated and miRNAs differentially expressed between groups (LP-12d versus NP-12d and LP-16w versus NP-16w) were evaluated. miRNA data have been generated following the MIQE guidelines [11].

### 2.6 Target prediction

*In silico* target, the prediction was performed for differentially expressed miRNAs using the combined analysis of three algorithms based on conservation criteria TargetScan [44], microRNA.org [38] and PicTar [39]. Results were taken from each search analysis and cross-referenced across all the three research results. To exclude the hypertension effect on gene expression, only targets predicted in both 12-day and 16-week old animals were considered. Furthermore, only targets genes expressed in cardiac tissue were used for the analysis. To offer experimental support to *in silico* predicted targets, we evaluated the gene expression by RT-qPCR and quantified the protein levels by western blot analysis.

### 2.7 Heart LV Predicted Gene Expression

Total RNA was extracted from LV of eight offspring in each group using the Trizol method [15]. The total RNA quantity, purity and integrity was assessed as previously described for miRNAs expression analysis. For the cDNA synthesis, High Capacity cDNA reverse transcription kit (Life Technologies, USA) was used. For real-time PCR, 2 μl cDNA (40ng/ μl) was added to a master mix comprising 10 μl TaqMan^®^ Fast Advanced Master Mix (Life Technologies, EUA), 1 μl primer mix and 7 μl water for reaction. Water was used in place of cDNA as a non-template control. The cycling conditions were: 50°C for 2 minutes, 95°C for 20 seconds, 50 cycles of 95°C for 1 second and 60°C for 20 seconds. Amplification and detection were performed using the StepOne Plus (Life Technologies, EUA) and data acquired using the StepOne Software v2.1 (Life Technologies, EUA). Ct values were converted to relative expression values using the ΔΔCt method with offspring heart data normalized to GAPDH as a reference gene. IDT^®^ Integrated DNA Technologies provided primers for mRNA RT-qPCR.

### 2.8 Western blot analysis

Some animals in each group were used to perform the protein level analysis by western blot. Immunoreactive bands were detected using the chemiluminescence method, and the intensity was quantitated by optical densitometry (Scion Image Corporation).

### 2.9 Statistical Analysis

Data are expressed as the mean ± standard deviation or as the median with interquartile range [lower quartile - upper quartile] and was previously tested for normality and equality of variance. Comparisons between two groups were performed using Student’s t-test when data were normally distributed and the Mann-Whitney test when distributions were non-normal. Comparisons between two groups through the weeks were performed using 2-way ANOVA for repeated measurements test, in which the first factor was the protein content in pregnant dam’s diet and the second factor was time. When an interaction was found to be significant, the mean values were compared using Tukey’s post hoc analysis. Significant differences in miRNA expression were detected using a moderated t-test. Data analysis was performed with Sigma Plot v12.0 (SPSS Inc., Chicago, IL, USA). The significance level was 5%.

## 3. RESULTS

### 3.1 Effect of protein restriction on body weight and food intake of pregnant rats

An interaction between the factors protein content in pregnant dam’s diet and the time was found analyzing the body weight, the food and the protein intake of the pregnant dams. Dams on LP diet during pregnancy were lighter on second and third weeks of pregnancy compared with dams on NP diet, despite an equal body weight in the first week of pregnancy (p_diet x time_ < 0.001; p_diet_ = 0.007; p_time_ < 0.001; Figure 1A). Thus, considering the entire pregnancy weeks, dams in the LP group had lower weight gain than those in the NP group (NP (n=21): 109.86 ± 20.04 g; LP (n=31): 87.00 ± 14.01 g; p<0.001).

**Figure 1.**
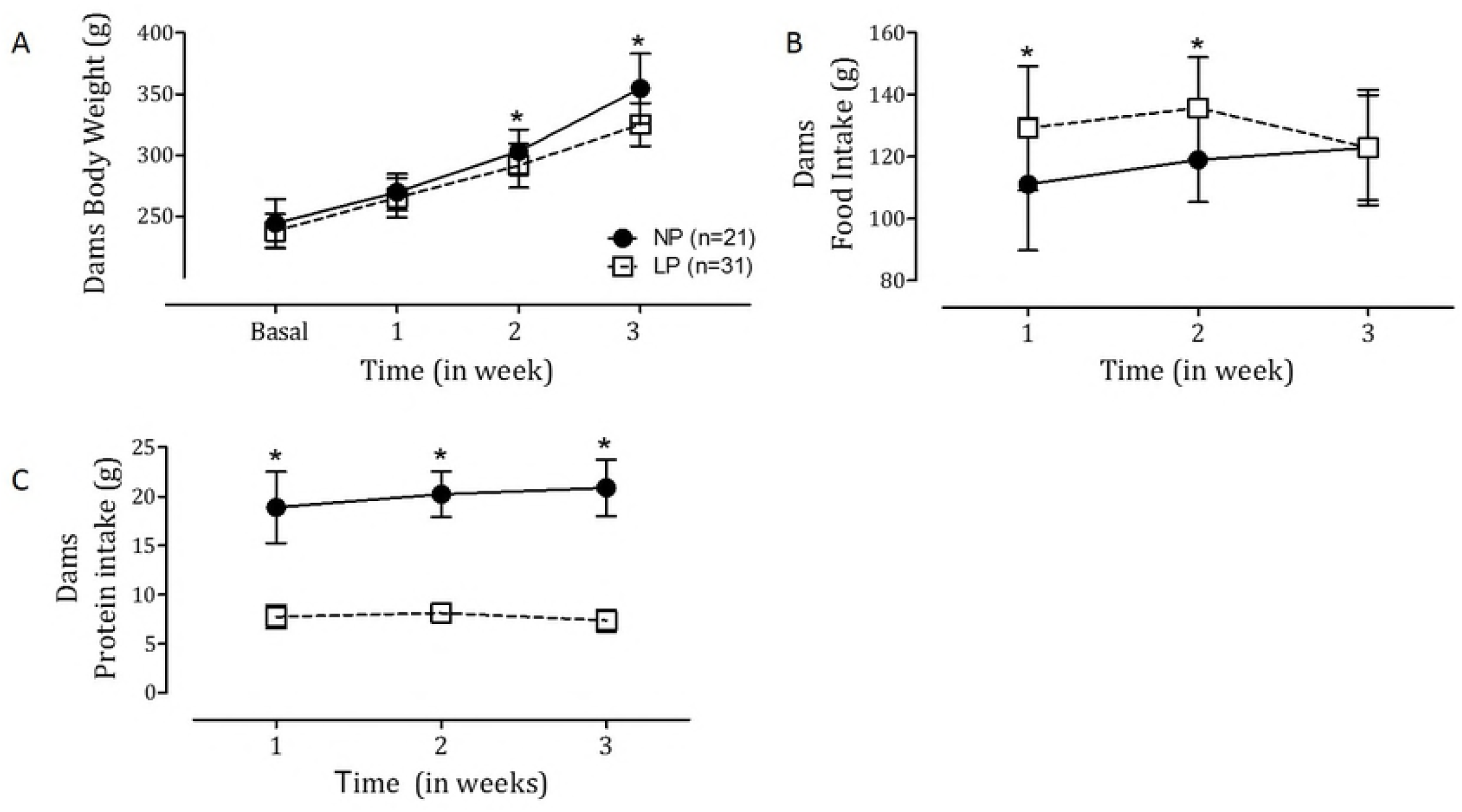
Body weight, food and protein intake of pregnant dams during gestation. Data were expressed as the mean ± standard deviation and was previously tested for normality and equality of variance. Comparisons between two groups through the weeks were performed using 2-way ANOVA for repeated measurements test, in which the first factor was the protein content in pregnant dam’s diet and the second factor was time. When an interaction was found to be significant, the mean values were compared using Tukey’s post hoc analysis. NP: normal protein diet group; LP: low protein diet group. A: Weekly body weight P_interaction_<0.001; P_diet_=0.007; P_time_<0.001. B: Weekly food intake. P_interaction_=0.018; P_diet_<0.001; P_time_=0.118. C: Weekly protein intake. P_interaction_=0.018; P_diet_<0.001; P_time_=0.069. Asterisk indicates a significant difference between week-matched NP (n=21) x LP (n=31) groups. The significance level was 5% *vs*. NP.

The weekly food intake was higher in LP dams in the first two weeks of pregnancy compared to NP dams (p_diet x time_ = 0.018; p_diet_ < 0.001; p_time_ < 0.118; Figure 1B). However, in the last week of pregnancy, there was no difference in food intake between the groups. Despite the higher food intake by LP dams during pregnancy, the assessment of weekly protein intake showed that dams from the LP group were exposed to severe protein restriction during the entire pregnancy (p_diet x time_ = 0.018; p_diet_ < 0.001; p_time_ = 0.069; Figure 1C).

### 3.2 Effect of gestational protein restriction on offspring phenotype

Male offspring from LP dams had lower birth weight (NP: 6.3g [6.0-6.7]; LP: 5.3g [4.9-5.9]; p<0.001) and higher anogenital distance (NP (n=103): 1.24 mm [1.15-1.45]; LP (n=107): 2.35 mm [1.24-1.49]; p=0.018) compared with offspring from NP dams. At 12 days after birth, offspring from NP-12d and LP-12d groups showed no significant difference on body weight (NP (n=11): 24.30±2.30 LP (n=13): 22.56±2.96; p=0.126).

Measurements on offspring in the 16-week groups showed no interaction between the factors and no significant difference related to protein content in pregnant dam’s diet were observed for weekly body weight (p_diet x time_ = 0.223; p_diet_ = 0.173; p_time_ < 0.001; Figure 2A). Analyzing the weekly food intake, although there was no interaction effect, the protein content in pregnant dam’s diet influenced this variable and, in general, LP offspring had lower food intake than NP offspring (p_diet x time_ = 0.275; p_diet_ < 0.001; p_time_ < 0.001; Figure 2B).

**Figure 2.**
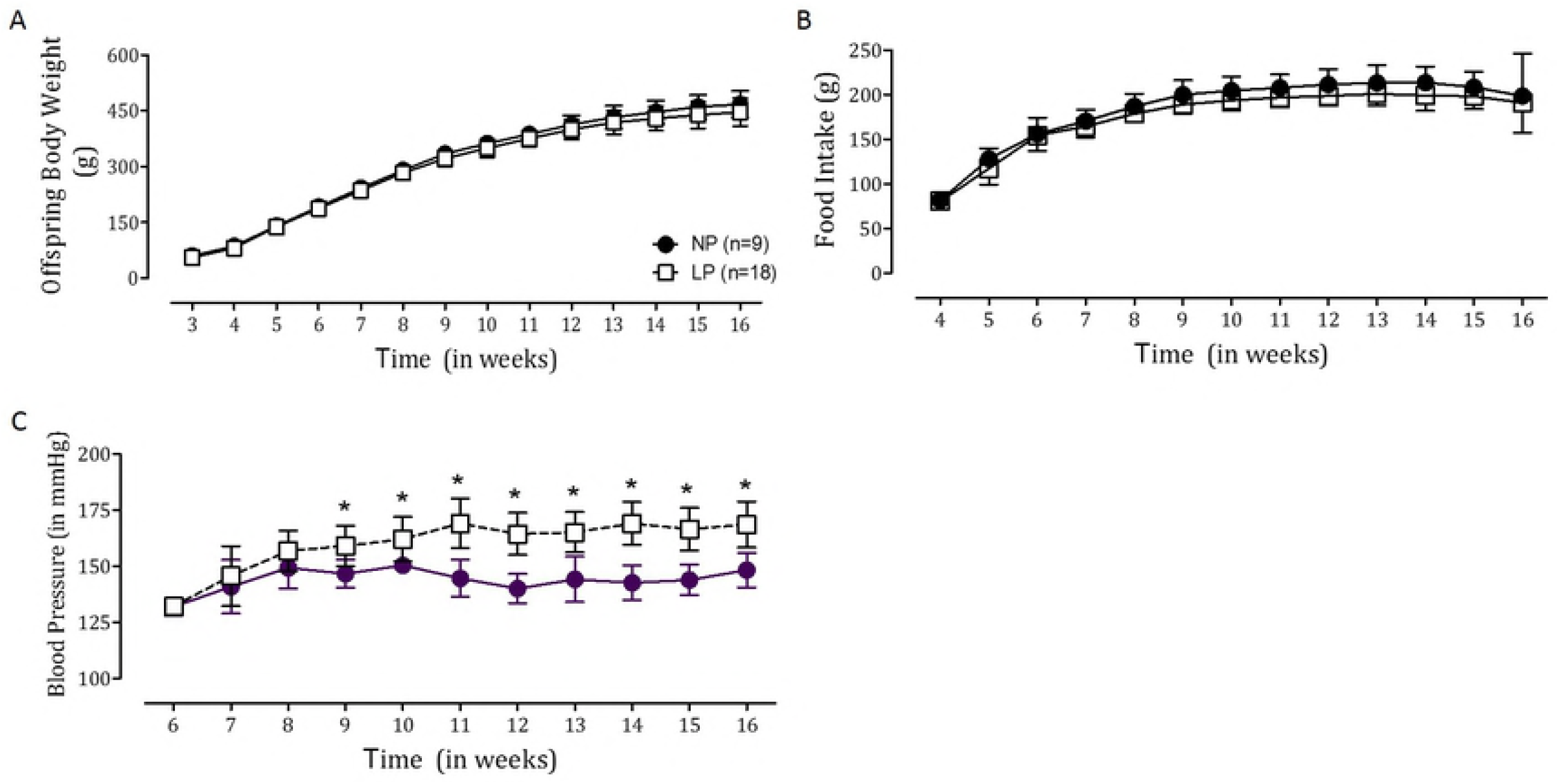
Body weight, food and systolic blood pressure of the 16-week old groups. Data were expressed as mean ± SD and was previously tested for normality and equality of variance. Comparisons between two groups through the weeks were performed using 2-way ANOVA for repeated measurements test, in which the first factor was the protein content in pregnant dam’s diet and the second factor was time. When an interaction was found to be significant, the mean values were compared using Tukey’s pos1 hoc analysis. NP-16w (n=9): normal protein diet group followed until 16 weeks old; LP-16w (n=18): low protein diet group followed until 16 weeks old. A: Weekly body weight. P_interaction_=0.223; P_diet_=0.173; P_time_<0.001. B: Weekly food intake. P_interaction_=0.275; P_diet_<0.001; P_time_<0.001. C: Systolic blood pressure. P_interaction_<0.001; P_diet_<0.001; P_time_<0.001. Asterisk indicates a significant difference between week-matched NP-16w x LP-16w groups. *P ≤ 0.05 *vs*. NP.

An interaction between the factors protein content in pregnant dam’s diet and time was found analyzing the systolic blood pressure from 6 to 16 weeks after birth. Animals from the LP-16w group had higher systolic blood pressure during 9 to 16 weeks of age compared to age-matched NP-16w group (p_diet x time_ < 0.001; p_diet_ < 0.001; p_time_ < 0.001; Figure 2C).

The morphometric analysis of the heart showed no differences were found for the LV weight comparing NP-12d versus LP-12d groups for both normalization to body weight (p=0.453) and tibia length (p=0.337) and then comparing NP-16w versus LP-16w groups for both normalization to body weight (p=0.796) and tibia length (p=0.259) (Table 1). Otherwise, the histologic analysis of the heart showed that the LP-16w group had higher myocyte CSA (p<0.001) and higher interstitial collagen volume fraction (p<0.001) compared to the NP-16w group (Table 1; Supplementary Figure 1).

**Table 1.** Cardiac left ventricle weight, myocyte cross-sectional area and interstitial collagen volume fraction. Data are expressed as the mean ± SD and was previously tested for normality and equality of variance. Comparisons between two age-matched groups were performed using Student’t-test. NP-12d: normal protein group followed until 12 days old; LP-12d: low protein group followed until 12 days old. NP-16w: normal protein group followed until 16 weeks old; LP-16w: low protein group followed until 16 weeks old. LV: left ventricle; CSA: myocyte cross-sectional area. *P ≤ 0.05 *vs*. NP.

|  | NP-12d | LP-12d | p | NP-16w | LP-16w | p |
| --- | --- | --- | --- | --- | --- | --- |
| **LV weight/**<br>**Body weight**<br>**(kg/g)** | 3.11±0.42(n=6) | 2.97±0.15(n=6) | 0.453 | 1.80±0.10(n=8) | 1.81±0.12(n=13) | 0.796 |
| **LV weight/**<br>**Tibial length**<br>**(kg/mm)** | 6.36±0.63 (n=6) | 5.88±0.75 (n=3) | 0.337 | 20.48±1.72 (n=5) | 19.38±1.73 (n=11) | 0.259 |
| **CSA (μm<sup>2</sup>)** | 27.4±2.7 (n=10) | 27.6±0.9 (n=11) | 0.777 | 139.0±6.2 (n=10) | 212.5±14.5 (n=13) | <0.001 |
| **Interstitial**<br>**collagen**<br>**fraction (%)** | 0.94±0.17 (n=9) | 1.00±0.22 (n=11) | 0.553 | 1.24±0.26 (n=10) | 2.26±0.55 (n=13) | <0.001 |

Data are expressed as the mean±SD and was previously tested for normality and equality variance. Comparisons between age-matched groups were performed using Student's t-test. NP-12d: normal protein group followed until 12 days old; LP-12d: low protein group followed until 12 days old. NP-16w: normal protein group followed until 16 weeks old; LP-16w: low protein group followed until 16 weeks old. LV: left ventricle; CSA: myocyte cross-sectional area. \*P ≤ 0.05 vs. LP.

### 3.3 Effect of gestational protein restriction on offspring miRNA expression in early-life and adulthood

Regarding heart, left ventricle miRNA expression, protein-restricted diet during pregnancy was significantly associated with male offspring altered miRNAs expression in both early life and adulthood. LP-12d versus NP-12d miRNAs fold-change depicted by volcano-plot showed a significant change in miRNA expression in early life (Supplementary Figure 2A). LP-12d group was associated with significant up-regulation of mir-184 (p=0,007), mir-192 (p=0,019), mir-376c (p=0,029) mir-380-3p (p=0,029), mir-380-5p (p=0,028), mir-451 (p=0,013) and mir-582-3p (p=0,029) and significant down-regulation of mir-547 (p=0,022) and mir-743a (p=0,004) compared to NP-12d group (Figure 3A). Volcano plot data from LP-16w versus NP-16w depicted in Supplementary Figure 2B shows a significant change in miRNA expression in adulthood. The LP-16w group had significant up-regulation of let-7b (p=0.017), mir-125a-3p (p<0.001), mir-142-3p (p=0.035), mir-182 (p=0.025) and mir-188-5p (p=0.029) and significant down-regulation of let-7g (p=0.045), mir-107 (p=0.021), mir-127 (p=0.029), mir-181a (p=0.045), mir-181c (p=0.029), mir-184 (p=0.029), mir-324-5p (p=0.006), mir-383 (p=0.002), mir-423-5p (p=0.006) and mir-484 (p=0,034) when compared to NP-16w group (Figure 3B).

**Figure 3.**
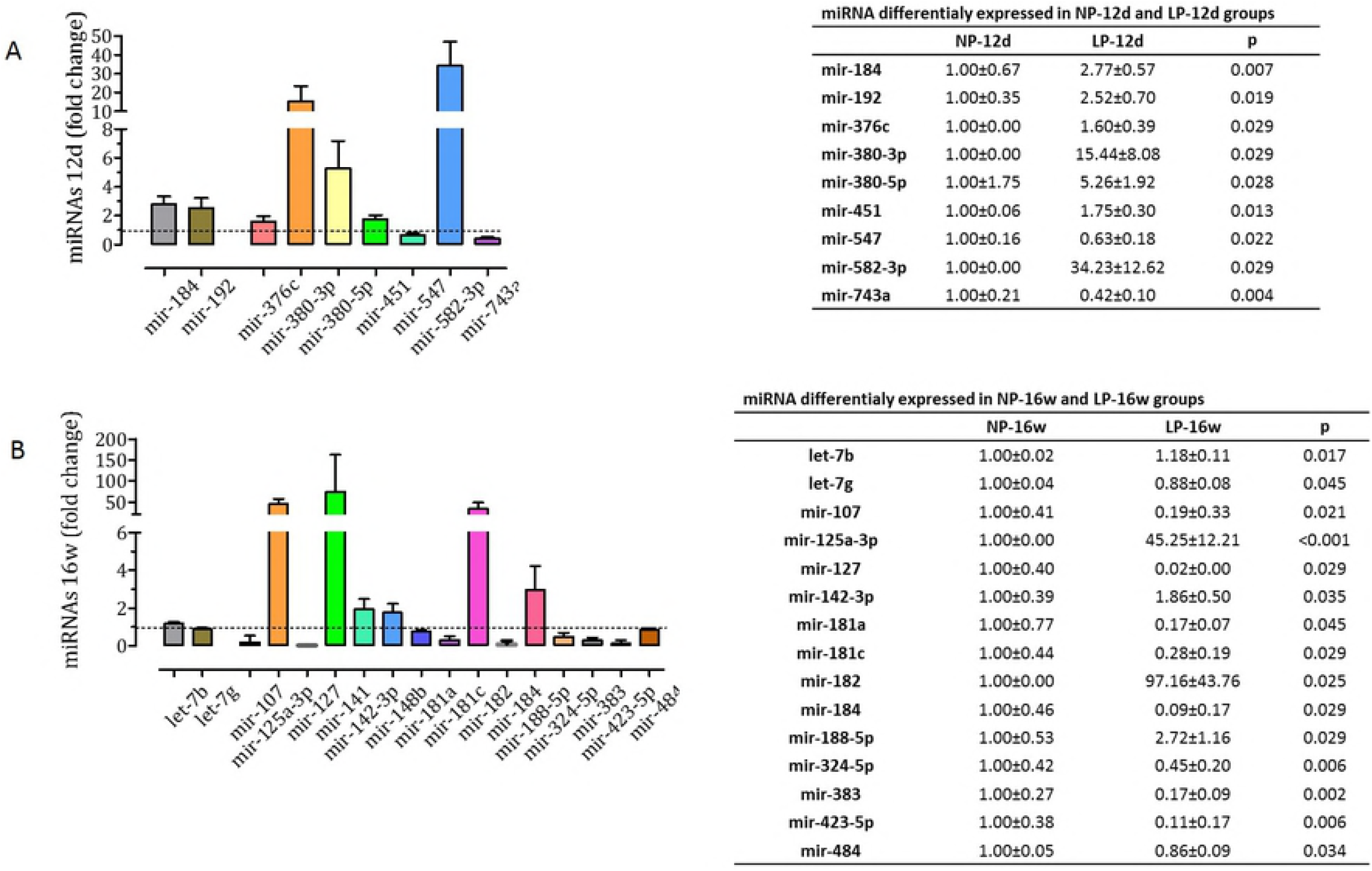
Differentially expressed miRNA of 12 days and 16 weeks old animals. A: Fold-change and miRNA expression values in LP-12d (n=4) versus NP-12d (n=4); B: Fold-change and miRNA expression values in LP-16w (n=4) versus NP-16w (n=4). Asterisk indicates a significant difference between age-matched groups. Significant differences in miRNA expression in age-matched groups were detected using a moderated Student’s t-test. *P ≤ 0.05 *vs*. NP.

### 3.4 Target prediction analysis

To test for potential mRNA targets of differentially expressed miRNAs, computational mRNA target prediction was performed. Many target mRNAs were identified for each of the miRNAs, although the number of targets varied per miRNA. By exploring the targets of the miRNAs by computational prediction, we had 165 possible mRNA targets for differentially expressed miRNA in the LP-12d group, and 281 mRNA targets for differentially expressed miRNA in the LP-16w group. Confronting the predicted mRNAs for both 12-day and 16-week old groups, we had 54 possible targets predicted simultaneously for both groups. Furthermore, selecting only the mRNA targets expressed in heart, we had 24 possible mRNA targets. Table 2 shows all the 24 mRNA targets considered for the analysis and the respective regulatory miRNA.

**Table 2.** Predicted mRNAs for the differentially expressed miRNAs in NP-12d versus LP-12d and NP-16w versus LP-16w groups.

| Gene symbol | Gene Name | Differentially expressed miRNA that could regulate the gene |  |
| --- | --- | --- | --- |
|  |  | NP-12d/ LP-12d | NP-16s/ LP-16s |
| **Adrbk1** | *Adrenergic receptor kinase, beta 1* |  | mir-423-5p |
| **Akap12** | *A kinase (PRKA) anchor protein (gravin) 12* | mir-184 | mir-184 |
| **Amotl1** | *Angiomotin-like 1* | mir-184 | mir-184 |
| **Bbs1** | *Bardet-Biedl syndrome 1 (human)* | mir-184 | mir-184 |
| **Calml3** | *Calmodulin-like 3* | mir-743a |  |
| **Dab2** | *Disabled 2, mitogen-responsive phosphopro.* | mir-743a |  |
| **Dnmt3a** | *DNA methyltransferase 3A* | mir-582-3p | mir-423-5p<br>mir-484 |
| **Gpr22** | *G protein-coupled receptor 22* | mir-192 | mir-182 |
| **Hbegf** | *Heparin-binding EGF-like growth factor* | mir-376c | mir-127 |
| **Hic2** | *Hypermethylated in cancer 2* | mir-547 | mir-127<br>mir-125a-3p<br>mir-484 |
| **Inpp1l** | *Inositol polyphosphate phosphatase-like 1* | mir-184 | mir-184 |
| **Insr** | *Insulin receptor* | mir-743a<br>mir-582-3p |  |
| **Jcad** | *RIKEN cDNA 9430020K01 gene* | mir-743a | mir-423-5p |
| **Mcf2l** | *Mcf.2 transforming sequence-like* | mir-184 | mir-184 |
| **Mmp8** | *Matrix metalloproteinase 8* | mir-184 | mir-184 |
| **Nfat5** | *Nuclear factor of activated T cells 5* | mir-380-5p | mir-324-5p |
| **Odc1** | *Ornithine decarboxylase, structural 1* | mir-743a | mir-423-5p |
| **Oxct1** | *3-oxoacid CoA transferase 1* | mir-743a | mir-324-5p |
| **Ppp2ca** | *Protein phosphatase 2, catalytic subunit* | mir-547 | mir-141 |
| **Rictor** | *RPTOR independent companion of MTOR, complex 2* | mir-192 | let7g<br>mir-142-3p<br>mir-188-5p |
| **Sirt1** | *Sirtuin 1* |  | mir-141 |
| **Tgfbr1** | *Transforming growth factor, beta receptor I* |  | mir-125a-3p |
| **Trps1** | *Trichorhinophalangeal syndrome I* | mir-547 | mir-484 |
| **Ubn1** | *Ubinuclein 1* | mir-184 | mir-184 |

### 3.5 Experimental support for predicted regulatory targets: Target’s mRNA RT-qPCR analysis

miRNAs can regulate post-transcriptional gene expression by targeting mRNAs for degradation. To explore the potential extent of miRNA-directed regulation of mRNA levels, RT-qPCR was used to measure mRNAs predicted to be targeted by the differentially expressed miRNA. The sequences of the primers used are shown in Supplementary Table 1. The results of the RT-qPCR analysis showed that the expression of Bbs1 and Calml3 genes were downregulated and that the expression of Dnmt3a, Oxct1, Rictor and Trps1 genes were upregulated in LP-12d versus NP-12d animals. Furthermore, the expression of Adrbk1, Bbs1, Dnmt3a, Gpr22, Inppl1, and Oxct1 genes were upregulated in LP-16w versus NP-16w animals (Figure 4; Supplementary Table 2). The expression of the other analyzed genes did not differ between groups.

**Figure 4.**
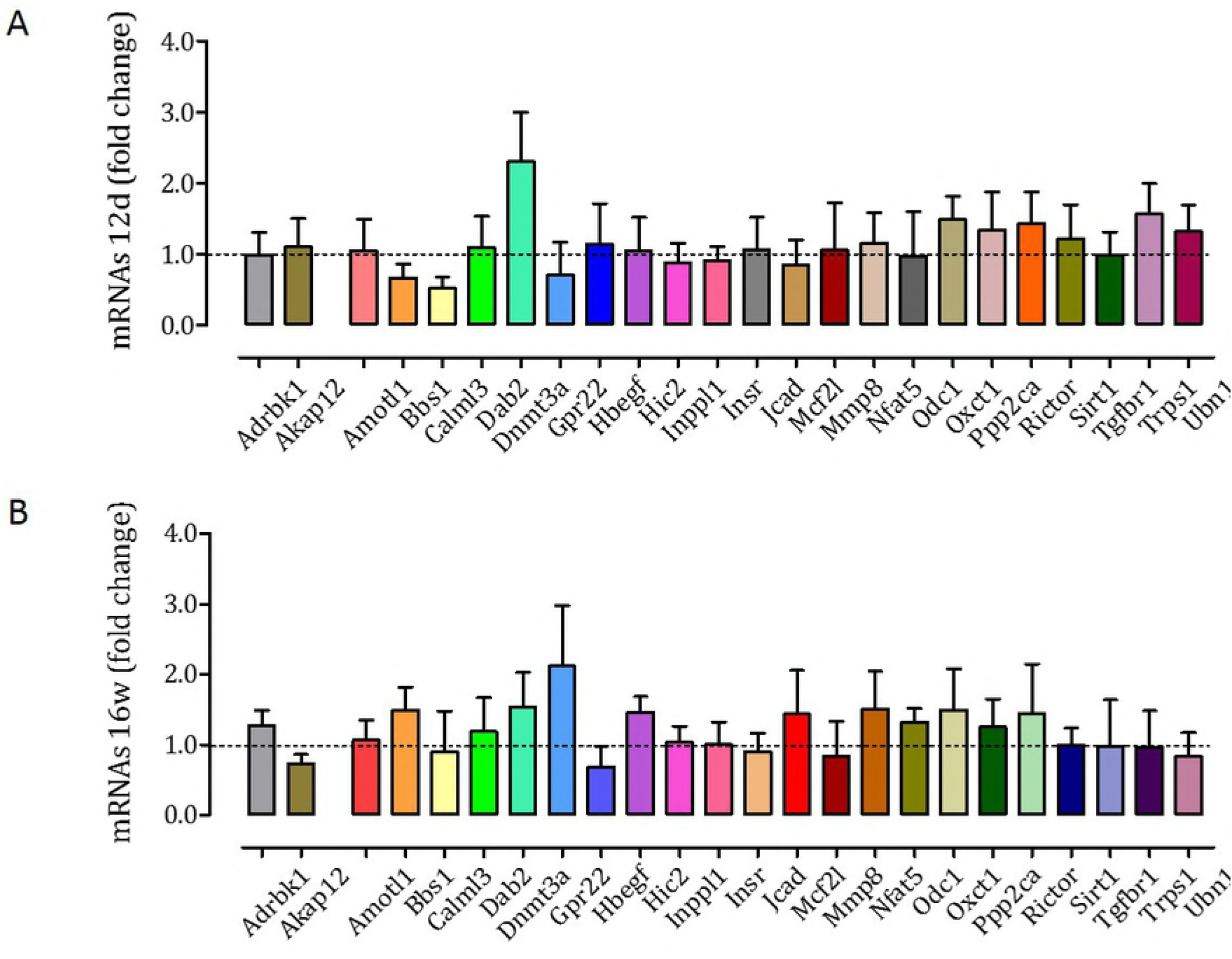
Targets mRNA expression of 12 days and 16 weeks old animals. A: Fold-change of mRNA expression in LP-12d (n=8) versus NP-12d (n=80; B: Fold-change of mRNA expression in LP-16w (n=8) versus NP-16w (n=8). Asterisk indicates a significant difference between age-matched groups. Significant differences in miRNA expression in age-matched groups were detected using Student’s t-test. *P ≤ 0.05 *vs*. LP.

### 3.6 Experimental support for predicted regulatory targets using western blot analysis

We quantified encoded proteins of genes whose expression was changed in the 12d groups thereby to exclude the possible effect of hypertension in the modulation of gene expression. The results of western blot analysis showed that LP-12d animals had lower levels of Bbs1 (NP-12d: 100.0±1.3; LP-12d: 94.7±1.8; p=0.027) and Calml3 (NP-12d: 100.0±4.33; LP-12d: 84.3±4.1; p=0.019) proteins in LV than NP-12d animals. Dnmt3a (NP-12d: 100.0±4.6; LP-12d: 141.3±11.5; p=0.017) and Oxct1 (NP-12d: 100.0±0.9; LP-12d: 112.3±3.0; p=0.037) proteins levels in LV were higher in LP-12d than NP-12d animals. No significant difference between these groups was found for Rictor (NP-12d: 100.0±2.6; LP-12d: 106.4±3.9; p=0.176) (Figure 5). The LP-16w animals had higher levels of Bbs1 (NP-16w: 100.3±3.4; LP-16w: 111.6±3.1; p=0.002) and Oxct1 (NP-16w: 100.0±3.1; LP-16w: 133.3±12.7; p=0.037) proteins in LV compared to NP-16s animals. No significant difference between these groups was found for Calml3 (NP-16w: 100.0±3.0; LP-16w: 102.3±4.2; p=0.232), Dnmt3a (NP-16w: 100.0±5.2; LP-16w: 108.3±7.5; p=0.424) and Rictor (NP-16w: 100.0±5.1; LP-16w: 102.6±5.8; p=0.749) (Figure 5). Trps1 protein was not detected.

**Figure 5.**
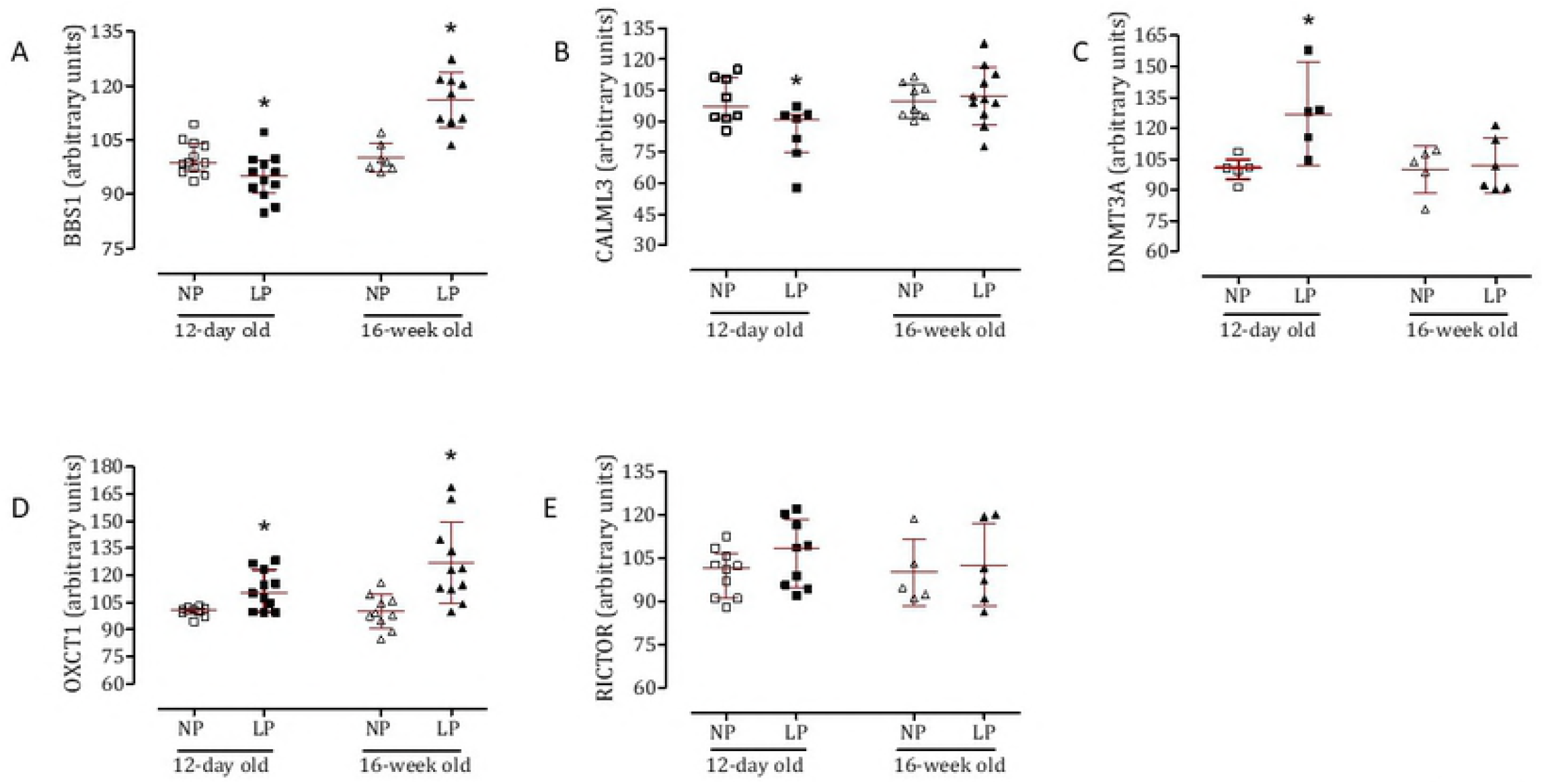
Western blot analysis of BBS1, Calml3, Dnmt3a, Oxct1 and Rictor of 12 days and 16 weeks old animals. Data are expressed as mean ± SD and was previously tested for normality and equality of variance. Comparisons between age-matched groups were performed using Student’t-test. *P ≤ 0.05 *vs*. LP. A: BBS1 in LP-12d (n=12) versus NP-12d (n=12) and LP-16w (n=12) versus NP-16w (n=9); B: Calml3 in LP-12d (n=13) versus NP-12d (n=8) and LP-16w (n=11) versus NP-16w (n=8); C: Dnmt3a in LP-12d (n=7) versus NP-12d (n=5) and LP-16w (n=7) versus NP-16w (n=5); D: Oxct1 in LP-12d (n=11) versus NP-12d (n=12) and LP-16w (n=15) versus NP-16w (n=11); E: Rictor in LP-12d (n=9) versus NP-12d (n=10) and LP-16w (n=6) versus NP-16w (n=5). P values of comparisons between age-matched groups were shown.

## 4. DISCUSSION

In the present study, protein-restricted offspring showed altered expression of a large number of heart LV miRNAs and predicted target gene expression was observed in both early life and adulthood LP offspring. Additionally, LP offspring had low birth weight, higher systolic blood pressure and changes in cardiac LV morphological parameters in adulthood compared with age-matched NP rats. Furthermore, this study showed that protein-restricted dams had a lower body mass gain and higher food consumption during pregnancy compared to NP rats. These results are supported by previous studies that showed the orexigenic stimulus and reduced mass gain on protein-restricted rats compared to isocaloric normal protein rats [20,27,73].

In protein-restricted chow, the carbohydrate content is approximately 15% higher than in normoproteic standard rodent chow. In this way, previous reports have shown that gastric emptying in LP animals is faster than in NP animals and consequently, the orexigenic signaling for food intake is rapidly triggered [27,73]. However, despite the increased food consumption, the experimental protein-restricted model proposed in the present study was kept.

In rats, the anogenital distance can be influenced by the embryo position in the womb, as well as by the sex of surrounding embryos due to the action of released steroid hormones [71]. Furthermore, anogenital distance is a sensitive marker of hormonal changes in rodents, especially from high and persistent steroid serum levels [34]. One of the mechanisms involved in the high exposure of the fetus to maternal glucocorticoids is related to a low concentration and decreased the activity of the placental enzyme 11beta-hydroxysteroid dehydrogenase (11β-HSD) type 2 [63]. Therefore, early fetal exposure to higher maternal glucocorticoids levels in gestational protein-restricted offspring may be responsible, at least in part, for the increased anogenital distance observed in protein-restricted offspring.

Gestational protein restriction is also associated with decreased intrauterine growth and low birth weight [17,20]. In the current study, we confirm these results, and we show that male offspring birth weight from protein-restricted dams was lower when compared to offspring from dams fed with a standard protein chow. The body mass was assessed from birth up to 16 weeks of life, and beyond the second week of age, we did not find a significant difference between LP and age-matched NP offspring. The recovery of offspring body mass after delivery in protein-restricted dams, known as “catch-up” is associated with a higher growth rate compared to the normal growth curve [78]. Furthermore, studies have demonstrated that fast mass gain after birth, in maternal LP offspring, is itself a risk factor for the development of hypertension [33], reduced peripheral insulin sensitivity and disorder in insulin secretion^61^, increased predisposition to obesity [53], metabolic syndrome [26] and increased cardiovascular risk [25]. Despite the occurrence of “catch-up” growth in 16-wk old LP offspring in the present study, this change was not related to increasing postnatal food intake when compared to age-matched NP animals.

16-wk old LP offspring showed an increased systolic blood pressure from the 9th week onwards when compared to age-matched NP animals. Several mechanisms may influence the development of hypertension in adults submitted to protein restriction during the intrauterine period. Clinical and experimental studies show that low birth weight due to both intrauterine growth restriction (IUGR) and maternal protein-restricted diet, are related to the reduction in the number of nephrons [7,50]. This kidney change, in turn, may alter glomerular hyperflow/hyperfiltration. Renal hyperperfusion/hyperfiltration accelerates glomerulosclerosis that naturally occurs with aging. The early loss of functional kidney units feeds a vicious cycle that perpetuates itself and determines the progressive increased renal retention of sodium and water and, consequently, enhanced arterial blood pressure [43,48]. However, the mechanisms related to hypertension development due to maternal nutritional impairment are complex and multifactorial. Although the impairment of nephrogenesis was associated with the hypertensive framework, fetal overexposure to glucocorticoids is a crucial component of this process [40]. Furthermore, endothelial dysfunction and loss of modulatory function performed by the vascular endothelium appear to be another critical element to the etiology of hypertension [51].

Regarding the evaluation of heart LV morphological findings, the current study has not shown any change in a whole organ or LV weight in both 12-day and 16-wk old LP offspring. However, the cardiac histological analysis in 16-wk old gestational protein-restricted rats showed a striking increase in the myocyte cross-sectional area associated with interstitial collagen expression in the LV. The literature is controversial about the heart weight of gestational protein-restricted offspring. In the rodent model, lower heart weight is often reported [17,75]. Otherwise, both higher [37] and equal heart weight [20,45] have also been reported in rats. These discrepant results may be related to several factors such as different strains used, protein-restricted intake levels, a period of hypoprotein diet restriction as well as differences in postnatal growth and arterial pressure values of LP offspring [78].

Additionally, the present study confirms previous studies showing the higher collagen content in heart LV in gestational protein-restricted rats compared with age-matched NP offspring [45,49]. The higher accumulation of collagen in the LV may compromise the myocardium elasticity and could be associated with functional cardiac disorders in adulthood [72]. Several authors have suggested that fibrosis occurs by hemodynamic overload imposed by arterial hypertension development. Furthermore, higher apoptosis [14] and reduced cardiomyocyte number [17] may explain the cardiomyocyte hypertrophy accompanied by increased collagen deposition in the left ventricle in LP offspring.

Regarding the miRNAs expression analysis, this work has identified nine miRNAs differentially expressed in 12-day old LP compared to age-matched NP offspring (upregulated miRNAs: mir-184, mir-192, mir-376, mir-380-3p, mir-380-5p, mir-451, mir-582-3p and, downregulated miRNAs: mir-547, mir-743a) and fifteen differentially expressed miRNAs in 16-wk old LP (upregulated miRNAs: let-7b, mir-125a-3p, mir-182, mir-188-5p and, downregulated miRNAs: let-7g, mir-107, mir-127, mir-181a, mir-181c, mir-184, mir-324-5p, mir-383, mir-423-5p, mir-484) compared to age-matched NP rats. Identification and validation of miRNA targets are of fundamental importance to gain a comprehensive understanding of miRNA function on modulation of cardiac phenotype in the present animal model of gestational protein restriction.

Analyzing the 12-day LP group, we observed the translation modulation of the following mRNAs encoding proteins Bbs1, Calml3, Dnmt3a, Oxct1, Rictor and Trps1. Furthermore, analyzing the 16-wk old LP group, we observed the translation modulation of the following mRNAs encoding proteins Adrbk1, Bbs1, Dnmt3a, Gpr22, Inppl1, and Oxct1. Then, in an attempt to exclude the possible bias due to hypertension in the LP-16w group, we evaluated the levels of proteins encoded by the genes that had altered expression in an LP-12d group versus NP-12d group. Thus, we performed western blot analysis to quantitate the level of Bbs1, Calml3, Dnmt3a, Oxct1, Rictor and Trps1 proteins. Trps1 protein level was not detected. The LP-12d group had lower Bbs1 and Calml3 protein levels and higher Dnmt3a and Oxct1 protein levels compared to the NP-12d group. Rictor protein level was similar in both LP-12d and NP-12d groups. The lp-16w group had higher Bbs1 and Oxct1 protein levels compared to the NP-16w group. Calml3, Dnmt3a and Rictor proteins levels did not differ between LP-16w and NP-16w animals.

Thus, it is evident that for some mRNAs targets the result of expression and protein level analysis was different from that expected by the respective miRNA analysis. Despite the lower levels of miR-743a, the expression and the protein level of the predicted target Calml3 gene was surprisingly lower in LP-12d versus NP-12d group. Similarly, the higher expression and protein level of Dnmt3a in LP-12d versus NP-12d groups was contrary to the expected higher expression of miR-582-3p. Also, the higher expression of miR-192 in LP-12d group disagreed with the higher expression and with the unchanged Rictor protein level in LP-12d versus NP-12d group. Furthermore, the higher expression of miR-182, the higher expression of the predicted target Gpr22 gene was surprisingly higher in LP-16w versus NP-16w group.

Several factors may explain this discrepancy between the expected and the obtained results after the analysis of miRNAs and their predicted targets expression. First, although not widely applicable, studies have suggested that miRNAs could also act as positive regulators of transcription [69,76]. Furthermore, it is evident that even the best available algorithms fail to identify a significant number of miRNA-gene interactions^55^. miRNA target prediction currently manages to detect 60% of all available targets and to provide one valid target in approximately every three predicted targets [70]. Finally, miRNAs integrate a high complexity network of gene expression regulation, and they have the potential to regulate a large part of the transcriptome [77]. Thus, each miRNA could regulate the expression of several mRNAs targets, and the expression of each mRNA target could be potentially regulated by several miRNAs [68]. However, despite these discrepancies, it is clear that in both 12-day and 16-week old animals, gestational protein restriction induces differential miRNAs expression that seems to have a modulatory function on the expression of specific genes that has been associated to cardiac morphology, metabolism, and function.

The Adrbk1 protein, also known as Grk2, under normal conditions, acts together with β-arrestin to promote the desensitization, internalization and reduce expression of β-adrenergic receptors after catecholamine stimulus [74]. However, hypertension [36] and heart failure [65] are associated with increased expression and activity of Grk2, which are initially linked to the prevention of excessive β-adrenergic stimulation. However, with chronic stimulation, a vicious cycle begins, and increasingly high levels of Grk2 contribute to heart failure progression [74]. Furthermore, the Grk2 expression is related to insulin resistance and increased mitochondrial stress [29,35].

The Inppl1 gene encodes a Ship2 protein that acts as a negative regulator of the insulin signaling pathway, decreasing the insulin sensitivity due to inhibition of Glut4 translocation [16]. Also, the Ship2 function is related to the inactivation of the PI3K-Akt signaling pathway [22]. Furthermore, Ship2 acts directly as docking protein to cytoskeletal proteins, focal adhesion proteins, and receptors associated with phosphatase and tyrosine kinase proteins [23,54].

Bbs1 protein is a structural component of cilia basal body and features a well-characterized role in the ciliary formation, stability, and function^3^. The bbs1 expression is related to reduced expression of insulin and leptin receptor plasmatic [59,62]. Furthermore, Bbs1 gene mutation is associated with higher susceptibility to congenital cardiac defects [21], heart valves and atrioventricular canal defects, dextrocardia and dilated cardiomyopathy [24].

Dnmt3a protein is one component of the DNA methylation epigenetic mechanism and, together with Dnmt3b and Dnmt3l are responsible for the methylation pattern establishing genomic DNA during the initial embryogenesis [32]. DNA methylation dynamics are essential during cardiovascular development as well as in the progression of cardiovascular disease. Heart failure in mice was associated with altered DNA methylation pattern that resembles the newborn pattern [30]. In an infarction rat model, Dnmt3a expression is increased due to the lower expression of mir-29a and mir-30c, and these changes correlate with post-ischemic tissue remodeling [28]. Furthermore, increased Dnmt3a expression is associated with lower RASSF1A expression in cardiac fibroblasts inducing, thereby, cardiac fibrosis [64].

Oxct1 protein is a critical component of the ketone body’s metabolism [18]. Although fatty acids are the primary energy substrate for myocardium [66], ketone body metabolism is physiologically crucial during the neonatal period [31]. In new-born rodents, ketogenesis is related to reduced white adipose tissue and altered availability of substrates after birth, since, during lactation, the availability of lipids is greater than carbohydrates while in the intrauterine period, the opposite availability occurs [18,31]. Furthermore, the heart energy demand increases after birth [18]. Ketone body metabolism is also essential in heart failure [18], and the change in energy substrate after cardiac injury seems to be a protective role against cardiac injury and ventricular remodeling [58]. However, the understanding of the pathophysiology of this metabolic switch as well as the context in which these changes are adaptive or maladaptive is limited [19]. Hepatic ketogenesis is stimulated, and plasma levels of ketone bodies increase in a heart failure model proportional to increase in blood pressure, leading to a reduction in fatty acids oxidation and increase in ketone body oxidation during the progression of cardiomyopathy [19].

## CONCLUSION

Thus, we conclude that all morphological heart alterations that were observed here in the protein-restricted offspring could be, at least in part, due to changes in cardiac miRNA expression. Some of the miRNAs differentially expressed in gestational protein restriction modulate the expression of several genes whose function is associated with cardiac morphogenesis and morphology by regulating cell polarity, the cytoskeletal dynamics and intracellular trafficking, cell proliferation, and growth, extracellular matrix deposition, and apoptosis. Furthermore, some miRNAs differentially expressed in this experimental model modulate the expression of genes whose function is associated with cardiac metabolism and function in the cardiovascular system. Although several studies have determined a close relationship between abnormal miRNA expression and human cardiac functional disorders, as far as we know, our study is the first description of changes in miRNA expression caused by gestational protein restriction that may modulate heart structure in early life and cause disease onset in later life.

Authors’ contributions
HBA: Data curation, Investigation, Formal analysis, Methodology, Visualization, Writing-original draft; JARG: Formal analysis, Methodology, Visualization, Writing-review & editing; PAB: Conceptualization, Formal analysis, Funding acquisition, Methodology, Resources, Supervision, Visualization, Writing-original draft, Writing-review & editing

## Acknowledgments

The authors thank Dr. Tom Fleming - University of Southampton for the collaboration with the paper review.

## List of abbreviations

11β-HSD: 11beta-hydroxysteroid dehydrogenase
2-way ANOVA: two-way analysis of variance
Adrbk1 gene: Beta adrenergic receptor kinase 1 gene
Bbs1 gene: Bardet-Biedl Syndrome 1 gene
Calml3 gene: Calmodulin-like protein 3 gene
cDNA: complementary DNA
COBEA: Brazilian College of Animal Experimentation
CSA: cross-sectional area
DNA: deoxyribonucleic acid
Dnmt3a gene: DNA methyltransferase 3 alpha gene
GAPDH: glyceraldehyde 3-phosphate dehydrogenase
Glut4: type 4 glucose transporter
Gpr22 gene: G-protein coupled receptor 22 gene
HE: hematoxylin and eosin
IDT^®^: Integrated DNA Technologies
Inppl1 gene: Inositol Polyphosphate Phosphatase Like 1 gene
LP: low protein diet
LV: left ventricle
miRNAs or mir-or miR: –MicroRNAs (miR-mature form of the miRNA; mir-pre-miRNA)
mRNA: messenger RNA
NIH: National Institutes of Health
NP: normal protein diet
Oxct1 gene: 3-Oxoacid CoA-Transferase 1 gene
PI3K-Akt: phosphatidylinositol 3’-kinase (PI3K)-Akt signaling pathway
Rictor gene: Rapamycin-insensitive companion of mammalian target of rapamycin gene
RNA: ribonucleic acid
Ship2: SH2 domain containing inositol 5-phosphatase 2
Trps1 gene: Transcriptional Repressor GATA Binding 1 gene
UNICAMP–: State University of Campinas

